# Deep Machine Learning provides state-of-the-art performance in image-based plant phenotyping

**DOI:** 10.1101/053033

**Authors:** Michael P. Pound, Alexandra J. Burgess, Michael H. Wilson, Jonathan A. Atkinson, Marcus Griffiths, Aaron S. Jackson, Adrian Bulat, yorgos Tzimiropoulos, Darren M. Wells, Erik H. Murchie, Tony P. Pridmore, Andrew P. French

**Affiliations:** The School of Computer Science, University of Nottingham,UK; The School of Biosciences, University of Nottingham, UK; Centre for Plant Sciences, Faculty of Biological Sciences, University of Leeds, Leeds, UK

## Abstract

Deep learning is an emerging field that promises unparalleled results on many data analysis problems. We show the success offered by such techniques when applied to the challenging problem of image-based plant phenotyping, and demonstrate state-of-the-art results for root and shoot feature identification and localisation. We predict a paradigm shift in image-based phenotyping thanks to deep learning approaches.

## Introduction and Background

The large increase in available genomic information in plant biology has lead to a need for truly high-throughput phenotyping workflows to bridge the increasing genotype-phenotype gap. Image analysis has become a key component in these workflows^1^, where automated measurement and counting has allowed for increased throughput and unbiased, consistent measurement systems. Machine learning has proven to be one of the most flexible and powerful analysis techniques, with approaches such as Support Vector Machines^2^ and Random Forests^3^achieving the highest success rates to date. Whilst these techniques provide considerable success in many situations^4^, their performance is saturating and often falls short of capturing the final 10% of accuracy required for fully automated systems.

Before introducing deep learning, it is helpful to first consider traditional machine learning techniques applied to bioimage analysis. It is generally assumed that raw images will contain too much information for a machine learning approach to efficiently process. For this reason, much of the established research in this field involves pre-computation of domain-specific image *features*, hand-crafted, for example, to detect areas of high contrast such as types of edges and corners. This pre-processing is intended to capture enough information to represent classes of objects, but contain significantly fewer dimensions than the full set of original image pixels^4^. The output of this feature detection is passed into a classifier, where classes (here, phenotypes) can be efficiently separated. Crucially, the choice of features is left tothe designer, and is often limited to existing sets, popular in the literature. These hand-crafted features are not guaranteed to provide the subsequent learning algorithm with the optimal description of the data, which in turn will reduce its effectiveness. It is easy to accidentally limit the application of the algorithm to specific tasks; an approach that performs well in one task may fail to perform in a different task. There is, therefore, a motivation to produce more general learning approaches.

Early general approaches include the biologically-inspired Artificial Neural Networks (ANNs), which use a set of simulated neuron-like connections, and transfer inputs via a set of learnt functions to a series of outputs. These represent a set of activations propagating through a network structure, triggered by input data, and resulting in an output activation pattern. ANNs typically use three layers, one for input, a hidden internal layer, and an output layer. Modern deep learning approaches extend this concept, and may contain many additional layers of artificial neurons (hence the term *deep*), and with increased complexity bring significantly-increased discriminative power^5^. Cutting edge algorithms and computation hardware have bought the training time for such networks down to practical levels achievable in most labs. Convolutional Neural Networks (CNNs) specialize this representation further, replacing the neuron-layers with feature-detecting convolution layers (biologically-inspired by the organisation of the visual field^6^, before finishing with traditional ANN layers to perform classification (**Fig.1**). CNNs have been quickly adopted by the computer vision community, but have also been used successfully in the life sciences^7^.

**Figure 1:**
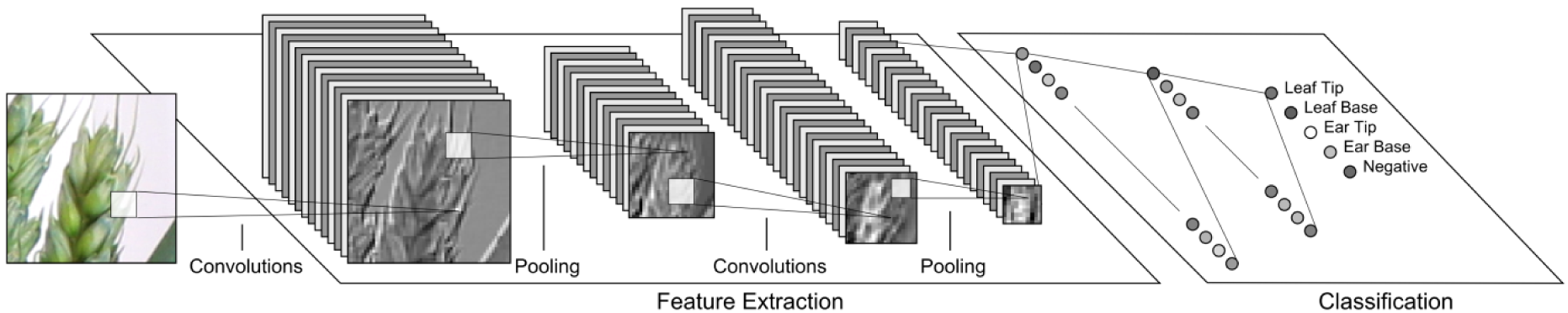
A simplified example of a CNN architecture operating on a fixed size image of part of an ear of wheat. The network performs alternating *convolution* and *pooling* operations (see online methods for details). Each convolutional layer automatically extracts useful features, such as edges or corners, outputting a number of feature maps. Pooling operations shrink the sizeof the feature maps to improve efficiency. The number of feature maps is increased deeper into the network to improve classification accuracy. Finally, standard neural network layers comprise the classification layers. which output probabilities for each class.

The CNN transforms feature maps from previous layers, creating a rich hierarchy of features that can be used for classification. For example, while the initial layer may compute simple primitives such as edges and corners, deeper into the network, feature maps based on these will highlight groups of corners and edges. Deeper still, feature maps may contain complex arrangements of features representing real-world objects^8^. These features are learnt by the CNN training algorithms, and are not hand-coded.

Modern CNNs will typically use many layers which makes training the networks complex, often requiring hundreds, sometimes thousands of images to train to the desired accuracy^9^. However,
once trained, their accuracy is unrivaled, and they can be transferred to other related domains by re-training using significantly fewer images^10^. A CNNis trained by iteratively passing example images containing the objects to be detected into the network, and adjusting the network parameters based on the results. The values of the convolutional filters are automatically adjusted to improve the result the next time a similar image is seen, a process that is repeated for as many images as possible.

To demonstrate the effectiveness of this deep learning approach, we trained two separate CNNs on two tasks central to plant phenotyping (rapid characterisation of plant physical and biological properties), framed as classification problems. In the first, given a small section of a root system image,can a CNN identify if a root tip present? The architecture of a root system is an important aspect of its physiological function; the root system’s structure allows it to access different nutrients and water within the soil profile. In phenotyping, particularly with high throughput 2D approaches, identifying features such as root tips represents the rate-limiting step in data quantification. We prepared training image data in which some images contained root tips, and some did not. This was derived from a dataset containing 2500 annotated images of whole root systems, and automatically generated classification images, by cropping at the annotated tip locations. This dataset will be made publically available.

**Figure 2:**
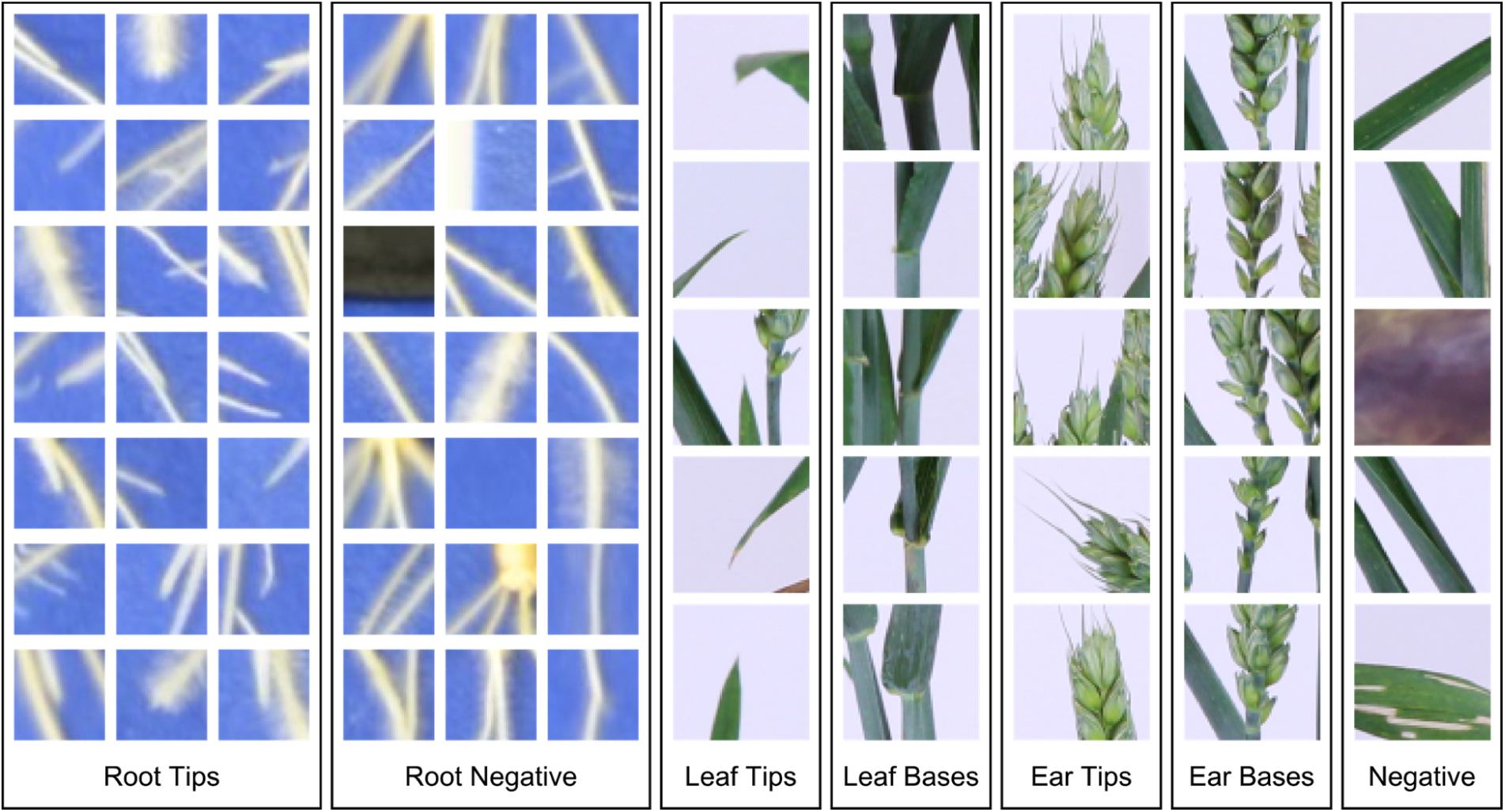
Example training and testing images from our root tip and shoot feature datasets. Positive samples were taken at locations annotated by a user. Negative samples were generated on the root system and at random for the root images, and on computed feature points on the shoot images.

In the second classification problem, given an image of a section of plant shoot, can a CNN identify biologically-relevant features such as leaf and ear tips, bases etc.? This would allow high-throughput phenotyping on an extremely large number of lines based on single images. It also allows 3D shotstructure to be linked with with physiological functioning: for example the separation into individual leaves and organs allows us to place biologically distinct plant parts within a useful functional context (different leaves, reproductive organs). To do this, we hand-annotated 1664 images of wheat plants, labelling leaf tips, leaf bases, ear tips, and ear bases. Classification images were then automatically extracted from these images as before. This dataset will also be made publically available. Example images from both datasets can be seen in (**Fig. 2**.

For both the root and shoot data, we randomly separated 80% of the data into a training set, and 20% remained for testing. To evaluate the accuracy of each network, we ran each test image through the network, obtaining the likelihood of each class. These were then compared to the true label for each image to ascertain whether the network had correctly classified the image. The accuracy of the root tip detection network was 98.4%. The shoot dataset, containing 4 classes of shoot features, along with numerous instances of cluttered, non-plant background, represents an even more challenging task. In this case, the shoot network successfully classified 97.3% of images. In both cases, CNNs here have out-performed recent state-of-the-art systems (e.g. accuracies of 80-90% have been typical^2^, ^11^). Accuracy results for individual classes can be seen in **Table 1**. Note also that both these scenarios are much more challenging than typical cell-scale successes seen to date,as the images involved are much less constrained.

As well as identifying features, it is necessary in quantitative phenotyping to locate the features within the image. For example, reliably identifying the locations of root tips is a bottleneck in automated root system analysis^12^, and is often omitted from image analysis software due tothe challenges localisation presents. Localisation of the different biological feature classes for a shoot is vital in capturing the architecture of the plant, essential for phenotyping. We have extended our root and shoot classifiers to perform localisation by scanning over each original image, applying the classifier to a window of the appropriate size centred on each selected pixel. As the output of the network is a set of class probabilities, pixels observed as above a likelihood threshold are marked as belonging to a specific class (**Fig.3**, **Supplementary Figure1**).

**Table 1:** **Classification results for both datasets. Leaf tips represent the hardest classificationproblem in the datasets, with large variations in orientation, size, shape, and colour. In all cases the accuracy has remained above 95%, with the average accuracy of both networks above 97%**.

| Feature | Correctly Classified | Misclassified | Accuracy (%) |
| --- | --- | --- | --- |
| Root Tip | 2904 | 73 | 97.5 |
| Root Negative | 5687 | 65 | 98.9 |
| Total/Average | 8591 | 138 | 98.4 |
| Leaf Tip | 2225 | 113 | 95.2 |
| Leaf Base | 2299 | 52 | 97.8 |
| Ear Tip | 686 | 15 | 97.9 |
| Ear Base | 765 | 23 | 97.1 |
| Shoot Negative | 6110 | 136 | 97.8 |
| Total/Average | 12085 | 339 | 97.3 |

**Figure 3:**
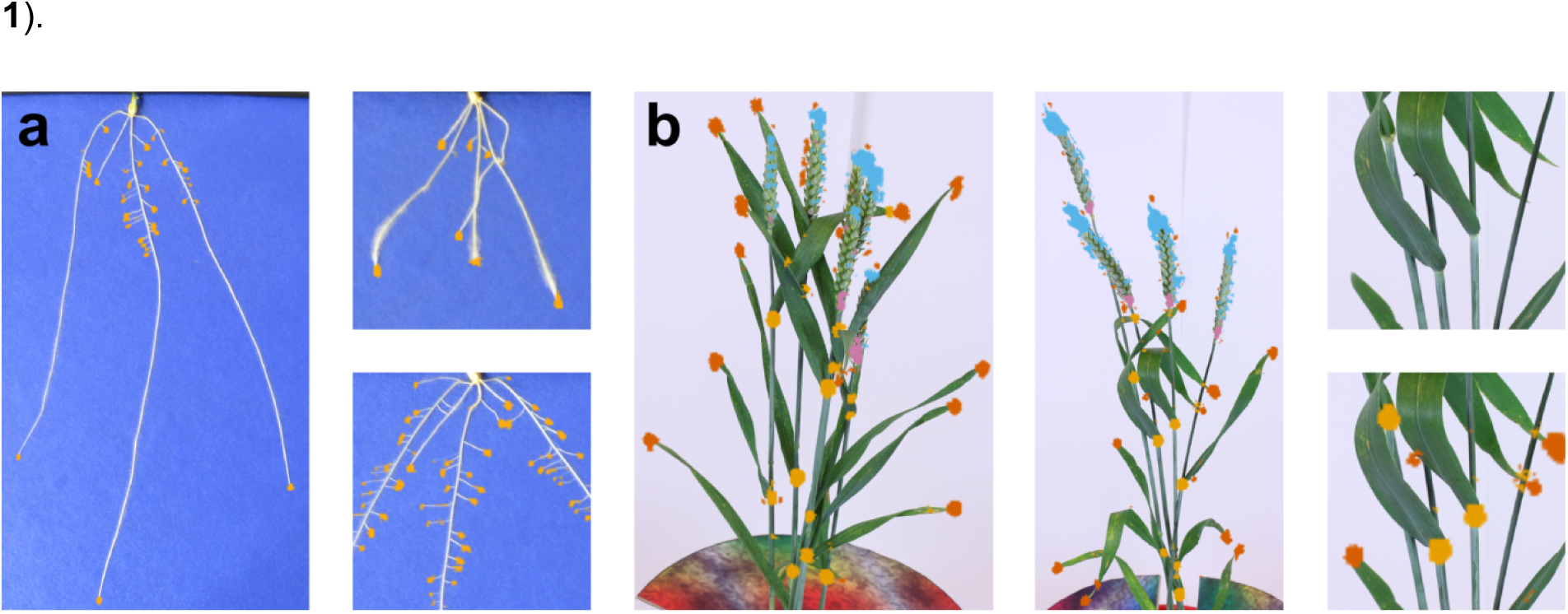
Localisation examples. Images showing the response of our classifier using a sliding window over each input image. (a) Three examples of wheat root tip localisation. Regions of high response from the classifier are shown in yellow. (b) Two examples of wheat shoot feature localisation. Regions of high response from the classifier for leaf tips are highlighted in orange, leaf bases in yellow, ear tips in blue, ear bases in pink. A portion of the second image has been zoomed and shown with and without features highlighted. More images can be seen in Supplementary Figure 1.

We have validated the accuracy of our localisation step by measuring the proportion of location windows containing false positives or negatives. The accuracy of the root tip location is 99.8%, the accuracy of the shoot feature location is 99.1% (**Supplementary Table 1**). Accuracy that is higher than that of the base classifiers presented earlier is not surprising. During training of the networks we generated particularly challenging negative examples of image features, these examples comprise only a very small fraction of each whole, real image.

## Conclusions

CNNs offer unparalleled discriminative performance in classification of images and localisation tasks. Here, we have demonstrated their efficacy of not only the classification, but also localisation of plant root and shoot features, significantly improving upon the state-of-the-art.

Deep learning is a very general technique, CNNs can be easily applied to other challenging problems, and determine useful features for classification automatically during training. Microscopy, x-ray, ultrasound, MRI or other forms of medicinal and structural imaging are all targets where deep learning will yield excellent results. We expect that the substantial increase in throughput offered by deep learning will lead to an improvement in the understanding of biological function akin to other high-throughput improvements in biology such as expression arrays^13^ and next-generation sequencing^14^, and anticipate numerous paradigm-shifting breakthroughs over the coming years.

## Online Methods

### Plant Growth and Imaging

#### Root Analysis

Winter wheat (*Triticum aestivum* L.) seedlings were grown and imaged as detailed previously^15^. After 9 days (two-leaf stage), individual pouches were transferred to a copy stand for imaging using a Nikon D5100 DSLR camera controlled using NKRemote software (Breeze Systems Ltd, Camberley, UK). Root system architectural traits were extracted from images of 2,697 seedlings using the *RootNav* software^12^ and used to produce the input images for CNN training.

#### Shoot Analysis

Wheat varieties were grown as detailed previously^16^. Plants were imaged according to the protocol of Pound et al.^17^. The developmental stage of each plants in both years of trial were the same. At anthesis, wheat plants (roots and shoots) were removed from the field and taken to a photography studio located close by to prevent wilting and damage to the shoots. They were imaged using three fixed Canon 650D cameras, with a minimum of 40 images per plant. Images were captured using a revolving turntable, including a fixed size calibration target. This target is used to facilitate 3D reconstruction, which does not feature in this work.

### Training and Testing Image Preparation

Convolutional Neural Networks (CNNs) using traditional neural network layers for classification can be applied to images of any reasonable size, but once trained at a certain size, this must remain consistent. We chose input sizes of 32x32 pixels for root tip images, and 64x64 pixels for shoot feature images. In the root case, a 32x32 image was found to be adequate to capture a root tip, along with enough information on the surrounding image. The 64x64 resolution of shoot features was chosen as a compromise between efficiency, and the higher resolution necessary to handle the more complex features seen in these images. Choosing a size appropriate to the feature of interest whilst maintaining a balance with computational efficiency is key here.

For root images, we obtained root tip positions from the database of annotated root systems, paired with the captured input images. For each source image, we cropped training images centred around each root tip position. This resulted in a variable number of training images per source image, depending on how many root tips had been annotated by the user. We restricted root tip images to primary and lateral roots that were longer than half the window size (16 pixels). Avoiding extremely short lateral root avoids ambiguity with root hair, which appears frequently on many of the images. For all training images in the root dataset, we cropped source images at 42x42 pixel size, and then performed an additional crop to 32x32 randomly during training. This approach, known as data augmentation, is akin to producing many more training images with random offsets, such that the root tips do not appear in the exact centre of each training image every time. This approach has been shown to produce improved accuracy when the classification target is not necessarily in the centre of each testing image, as maybe the case when we use our scanning localisation approach.

We also generated negative training images, which do not contain the features of interest, with two times more negative images than positive ones. We increased the number of negative images in order to adequately capture the wide variety of different negative images that are possible on in this data. Half of the negative data was generated at random points on the source image, but limited to areas that contained no root tips. The remaining negative data was generated at random positions on the known root system, again avoiding root tips. This is a form of hard negative mining, where negative testing data is generated on regions that appear similar to the positive data This has been shown to improve the accuracy of machine learning algorithms over negative data produced entirely at random^18^. The total number of images produced was 43,641, which was split at random into a training set of size 34,912 and a testing set of size 8,729.

We used a similar approach in the preparation of shoot feature images. For each source image we selected image crops at each annotated location. The shoot images are a higher resolution than the root images, we found that we obtained better accuracy if we cropped 128x128 images, then scaled to 64x64 for use in the network. This simply includes more of each image within the field of view of the smaller crops. We summed the number of each type of feature (e.g. leaf tip, ear tip) to produce an overall positive image count, and generated an equal number of negative images per source image. Unlike the root system data, where information on the position of the remaining root system could be used to generate hard negative data, the shoot annotation only included the specific features to be classified. In order to generate hard negative data, we used a Harris feature detector^19^ to generate candidate points of interest, then selected from this set at random. This ensured that the negative data contained large amounts of clutter and other plant material,rather than just background. Finally, we generated a small amount of additional images at random, to ensure that areas such as the white background were represented. The resulting dataset contained 62,118 images, of which 49,694 were training images, and the remaining 12,424 were used for testing.

### CNN Network Design

We used the Caffe deep learning library^20^ to develop each network. In Caffe, networks are described using a series of structured files, along with information on training and testing, such as how often to perform testing when training, and so-called hyperparameters, such as the learning rate, which will be described below.

We designed separate CNN architectures for each problem. These architectures are shown in (**Supplementary Fig. 2**); they adapt a common approach to CNNs, utilising multiple convolutional layers using 3x3 kernels, prior to each pooling layer^21^. The shoot CNN contains more layers to accommodate the larger input image size. It also includes increased feature counts in deeper layers, to address the more challenging classification task posed by the shoot images. Both networks end in neural network classification layers (often referred to as fully-connected layers) that reduce the output size to 2 and 5 respectively. Once trained, these final neurons represent the likelihood that the network has observed each class, and can be read to determine which class the network has identified.

The root CNN contained two groups of two convolutional layers, and one max pooling layer. Following these, two final convolutional layers perform further feature extraction, before three standard neural network layers performed the classification. The feature size of the convolutional layers was increased after each pooling layer, beginning at 64 convolutional filters, up to 256 filters. Finally, the neural network layers gradually reduce the feature size back down to 2, representing the separate “Root Tip” and “Root Negative” classes.

The shoot CNN contains three groups of convolutions and pooling layers. The number of convolutional layers between pooling layers varied slightly throughout the architecture in order to ensure that the spatial resolution of the data was always a multiple of two. A single final convolution is followed by three neural network layers performing the classification. The feature sizes of the convolutional and neural network layers were also increased over the root CNN. Feature sizes started at 64 filters, up to a maximum of 512 filters. The neural network layers decrease this feature size back down to 5, representing the 5 classes being detected.

Recent developments in CNNs have proposed additional components that improve performance. Neural networks require non-linear functions between layers in order to capture the complex non-linearity of the classification tasks. Traditionally, sigmoid or tanh functions have been used, where the result of each convolutional filter at each position is passed into a nonlinear function, before being passed to the next layer. Recent work^9^ proposed an alternative function, the non-rectified linear unit (“Relu”), which has been shown to improve the speed of training deep networks. We utilised Relu layers between all Convolutional layers, and between all fully-connected neural network layers. Other work^22^ proposed an approach whereby a percentage of fully-connected neurons are randomly deactivated during each iteration of training; this has been shown to avoid the overfitting problem, in which the classification of the training data improves, but at the expense of generality on the testing data. By deactivating neurons some of the time, the fully-connected layers are forced to learn from all parts of the network, rather than become focused on a few key neurons. We included dropout layers with a 50% dropout rate between the fully-connected layers.

### CNN Training and Validation

The Caffe library is built to perform iterative testing and training for as long as is required. Periodically the accuracy of the networks were measured using the separate testing data, and learning was halted after a steady state was reached, where no further improvement was seen if the network was left training. The learning rate specifies how quickly the network attempts to improve based upon the current set of images it is examining. This is an important feature of network learning; a low learning rate will mean the network does not adapt sufficiently fast to correctly classify images it sees. A learning rate that is too high may cause the network to wildly over-adapt, meaning it will improve on the current set of images, but at the expense of all images it has seen previously. As with most modern CNN approaches, we chose a higher learning rate to begin training, then periodically decreased this rate to “refine” the network to higher and higher accuracies. We began with a learning rate of 0.1, then decreased the learning rate by a factor of 10 every 20,000 iterations. In practice, we found that our networks were robust to changes in this learning rate, but that we stopped seeing any real improvement in accuracy when the learning rate fell below 1x10^−3^

Once training has completed, the learned parameters of the network are then stored and can be usedtoperform classification when required. The final accuracy of the networks described in this paper is the result of a final evaluation over all testing images once training was stopped. Our CNN models, learned parameters, and all the related scripts for training and testing will be made publically available.

### Localisation of Features

Determining where features appear in whole root or shoot images was performed by scanning the respective classifier over each image at regular pixel intervals (often referred to as a stride). Selection of the stride is straightforward, and is a compromise between pixel-wise accuracy of the resulting classification map, and computational efficiency. A stride of 1 will produce sub-images centred around every pixel, such that images will overlap the majority of the previous sub-image. This means that a feature visible in one image, will also be visible in a number of consecutive images around it. There is also significant computational cost to running a CNN network over this many sub-images, indeed such an approach is 4x slower than a stride of 2, and 16 times slower than a stride of 4 (as the stride applies in both the x-and y-axes). For both the root and shoot system images, we chose a stride of4, which results in a single scan taking under two minutes, and yet will output a classification map showing each feature location clearly. The scripts we used to perform this classification, and repeat this automatically over any number of images can be downloaded alongside our models.

For the root system images, we extracted sub-images at 32x32 resolution. To avoid truncation due to border effects, no images were centred around any pixels in the outer 16 pixels of the source image. Once scanning was complete, we upscaled the heatmap by a factor of 4 to return to the original image resolution. During scanning, the output from the network for each sub-image contains the likelihood that the image contains a root tip. We thresholded these values to produce a binary classification map,showing the locations of all detected tips. We found that a very strict threshold of 0.99 produced the best results; the network has been trained for an extended period, and we found that wherever a sub-image exhibited qualities that looked like a root tip, the likelihood output by the network was always extremely high.

For shoot images, we extracted sub-images at 64x64 pixel resolution. We began by reducing the sizeofthe original image by 50%, to account for our original image capture approach in which we cropped 128x128 pixel images. We then selected sub-images in an identical way to the root scenario, except that we extended the boundary at the edge of the input image to 32 pixels, to avoid truncation. The shoot CNN output 4 separate values for each sub-image, for the four features being detected. We thresholded these separately to produced a combined classification map for all features. We found that a threshold at or above 0.90 worked for all classes, however we fine tuned these thresholds slightly to achieve better classification accuracy. We found stricter thresholds e.g 0.99, were more effective in avoiding false-positives on leaf and ear tips. These parameters can be adjusted easily in our available scripts.

### Validation of Localisation

Validation was performed on 20 images for roots, and 20 for shoots. In both cases no images, or parts of these images, had been used in the training of either network. Accuracy was measured as the percentage of pixels that were correctly classified as either true-positives or true-negatives. False positives were determined as those pixels that were classified as a feature, but were outside of a radius around any ground truth features. This radius was set as half of the classification window size, in which any feature should be visible. False negatives were those pixels within the same radius of a ground truth feature that were not correctly classified as those features. Separate results for roots and shoots, and for each class, can be seen in (**Supplementary Table 1**). The scripts usedfor testing will be made available alongside our models.

---

**Supplementary Figure 1:**
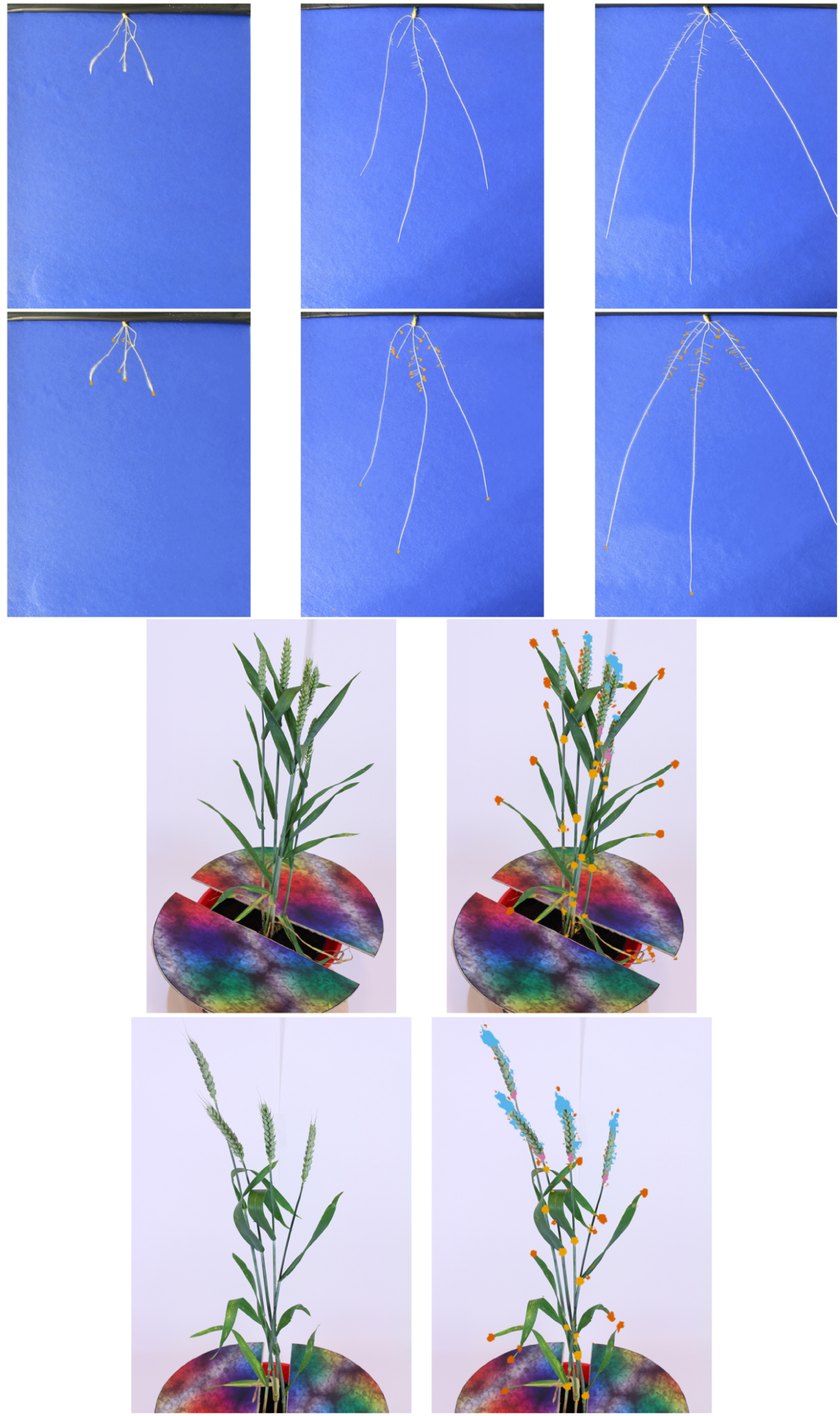
Original images showing the response of our classifier using a sliding window over each input image.

**Supplementary Figure 2:**
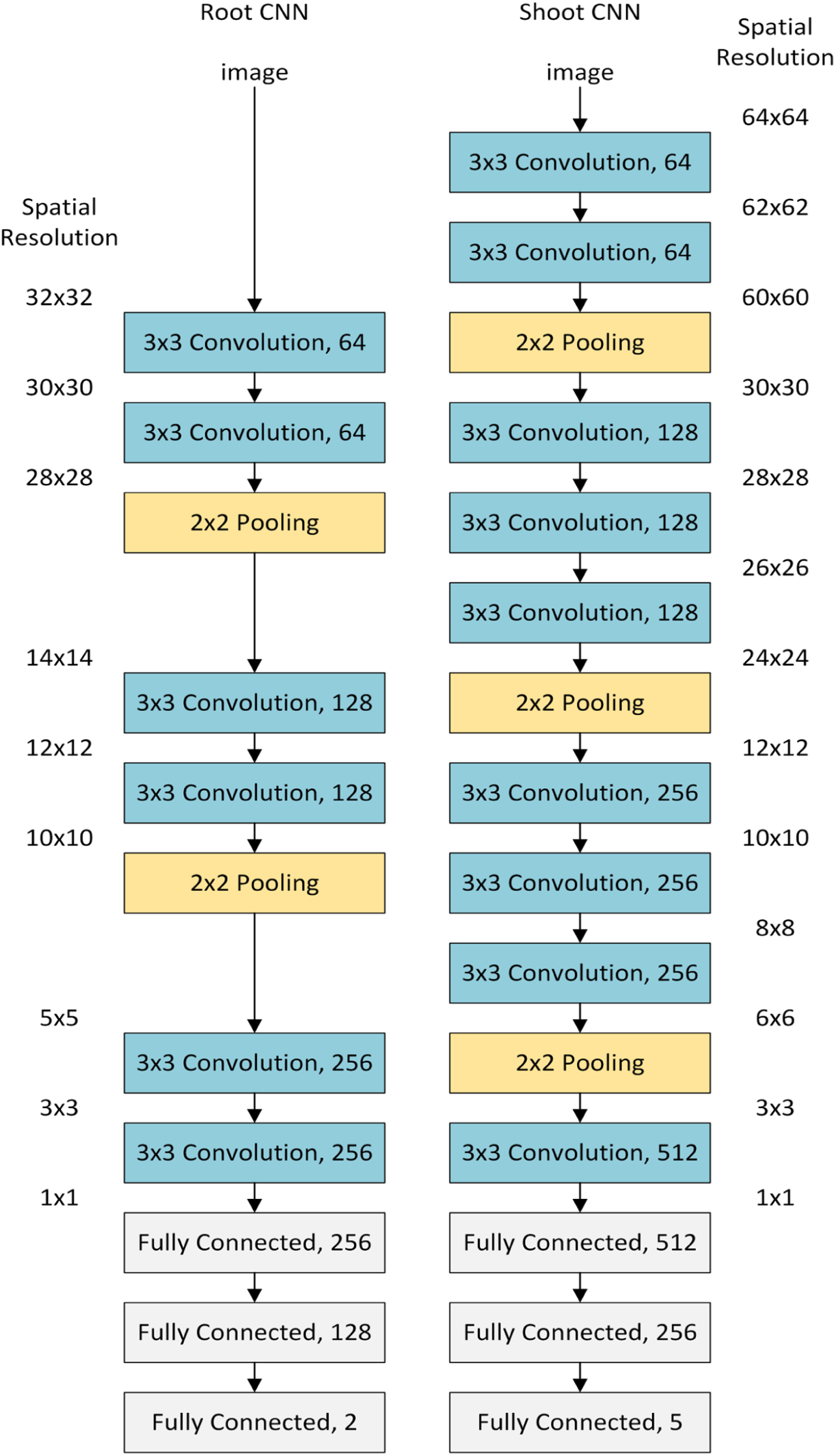
The architecture of both convolutional neural networks. In each case convolution and pooling layers reduce the spatial resolution to 1x1, while increasing the feature resolution. All convolutional layers used kernels of size 3x3 pixels, and the number of different filters is shown at the right of each layer. Following the convolution and pooling layers, the fully connected (neural network) layers perform classification of the images. We included ReLu layers between all convolutional and fully-connected layers, and dropout layers between each fully-connected layer.

**Supplementary Table 1:**
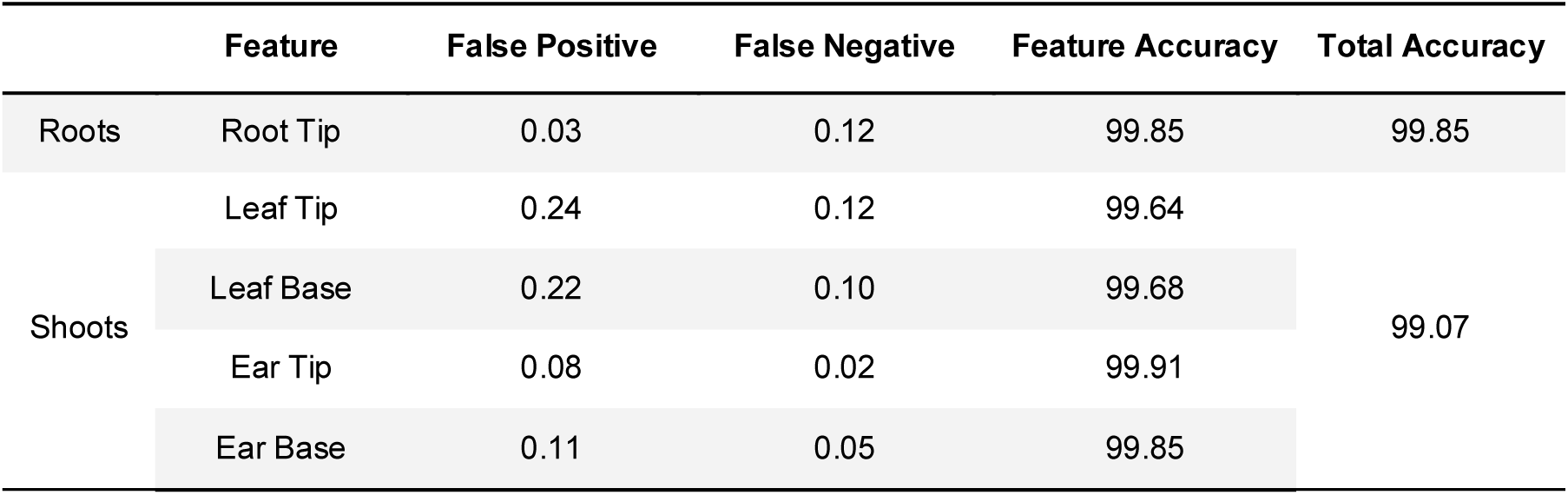
**Validation results for our image scanning approach over 20 root images, and 20 shoot images**.

